# A new bioorthogonal PET reporter gene for cardiac regenerative medicine binding to DTPA-lanthanide complexes

**DOI:** 10.64898/2026.09.27.754835

**Authors:** Sepideh Seyfi, Tatjana Dorn, Christine M. Poch, Martin Grashei, Eleonore Baier, Volker Morath, Veronika Fricke, Katja Fritschle, Arne Skerra, Markus Schwaiger, Christian Kupatt, Karl-Ludwig Laugwitz, Alessandra Moretti, Wolfgang A. Weber

## Abstract

The lack of non-invasive, quantitative methods to track the location and proliferation of transplanted cells hampers the clinical translation of cardiac regenerative therapies. Here, we establish a PET reporter for tracking human induced pluripotent stem cell (hiPSC)-derived cardiac cells across complementary two-dimensional (2D) and three-dimensional (3D) cardiac tissue models.

**Methods:** hiPSCs were genetically engineered using CRISPR-Cas9 to express a human-derived, anticalin-based, bioorthogonal PET reporter (DTPA-R) that specifically binds radiolabeled DTPA-metal complexes. The engineered hiPSCs were differentiated into ventricular progenitors (HVPs) and cardiomyocytes (CMs). Stability of reporter gene expression and its effects on cellular differentiation and function were evaluated. Uptake of [¹⁸F]F-DTPA•Tb by DTPA-R-positive and -negative HVPs was analyzed in vitro, and DTPA-R HVPs were evaluated in ex vivo cultured porcine myocardial slices with and without radiofrequency ablation (RFA) injury. Phantom studies were performed to evaluate PET detectability under clinically relevant large-animal and human imaging conditions and to assess how activity distribution affects signal detection in relation to DTPA-R cardiac cell-equivalent numbers.

**Results:** DTPA-R expression was maintained during the differentiation stages of hiPSC-derived cardiac cells and had no measurable impact on migration, differentiation, and functional integration. PET imaging enabled the specific detection of DTPA-R HVPs in intact and injured myocardial slices. At the initial phantom scan, the measured activity corresponded to cell-equivalent numbers of approximately 3.2×10⁶ HVPs or 5.5× 10⁶ CMs. The compact 200 µL source remained clearly detectable at 6 h, whereas the more dispersed 1,000 µL source progressively approached background activity, indicating that PET detectability depended strongly on local activity concentration and distribution.

**Conclusion:** This study establishes a bioorthogonal PET reporter gene platform for quantitative imaging of hiPSC-derived cardiac cells, with no measurable adverse effects on the biological parameters examined, and provides a methodological foundation for future translational and *in vivo* imaging studies. The approach may also be applicable to other hiPSC-derived cell products beyond cardiac cell lineages.

## Introduction

Heart failure remains a leading cause of morbidity and mortality, mainly due to the limited regenerative capacity of the adult human heart after injury (*1*). Cardiac injury typically results in loss of functional myocardium and fibrotic scar formation, limiting long-term functional recovery (*2*). However, cell-based therapies using human pluripotent stem cell (hPSC)-derived cardiomyocytes (CMs) or progenitors such as human ventricular progenitors (HVPs) have shown promise by integrating into the host myocardium and improving function in preclinical models (*3–5*). HVPs additionally exhibit properties particularly relevant to cardiac repair, including migration within injured myocardium and differentiation toward the CM lineage (*4*). Clinical translation of these approaches remains hampered by the lack of methods for quantitative, longitudinal, and non-invasive cell tracking.

Current strategies rely primarily on endpoint histology or direct cell-labeling. Direct cell-labeling suffers from signal dilution due to cellular proliferation and redistribution to phagocytic cells in case of cell death (*6–9*). Although histological and immunofluorescence analyses provide detailed information on cell identity and tissue integration, they do not permit longitudinal assessment within the same subject. Direct cell-labeling approaches for MRI, PET, and SPECT have therefore been explored; however, these approaches primarily track the label rather than the transplanted cells themselves (*6,10*).

Reporter gene imaging can overcome these limitations. By genetically engineering therapeutic cells to express a dedicated reporter, this strategy enables cell-specific and longitudinal detection beyond the persistence of an initially administered label (*11*). However, current reporter systems, such as the herpes simplex virus type 1 thymidine kinase (HSV1-tk) reporter used for PET imaging, raise immunogenicity concerns for long-term studies (*11–13*), whereas human-derived reporters such as the sodium/iodide symporter (NIS) can be limited by endogenous expression in thyroid and other tissues (*14*).

Additionally, reporter systems have rarely been validated in physiologically relevant cardiac environments, where mechanical and electromechanical integration are critical for the therapeutic success of the labeled cells.

We previously developed a PET reporter based on a membrane-anchored anticalin protein (DTPA-R) that enables high-affinity binding of the exogenous radiolabeled DTPA ligand [¹⁸F]F-DTPA•Tb. The reporter is expressed at the cell surface and can additionally be detected by antibody-based approaches, providing complementary imaging and tissue-level readouts. In chimeric antigen receptor (CAR) T cells, this reporter enabled quantitative and longitudinal PET tracking without measurably impairing T-cell function (*15*).

Here, we adapted the DTPA-R system for cardiac applications by genetically labeling human induced pluripotent stem cells (hiPSCs) and differentiating them into functional HVPs and CMs. We evaluated reporter stability, biological compatibility, and imaging performance in 2D and 3D cardiac models, including *ex vivo* myocardial tissue.

## Methods

### Generation of DTPA-R hiPSC reporter line

The DTPA-R gene was introduced into hiPSCs via targeted integration at the AAVS1 safe harbor locus **(Method S1)**. The DTPA-R plasmid encodes a cell-surface–anchored anticalin that binds CHX-A’’-DTPA-containing lanthanide complexes for PET imaging and a V5-tag for immunofluorescent detection (*15,16*). It was cloned under the control of the CAG promoter on an AAVS1 donor vector (Addgene #80488). Genome editing was performed using a CRISPR– Cas9-mediated homology-directed repair strategy adapted from established AAVS1 targeting protocols (*17*).

### DTPA-R hiPSC maintenance and cardiac differentiation

DTPA-R hiPSCs were maintained under feeder-free conditions on Geltrex-coated plates (Geltrex LDEV-Free; Gibco) in Essential 8 medium (E8; Gibco) at 37°C and 5% CO_2_, with daily medium changes. Cells were passaged every 3-4 days using 0.5 mM EDTA and supplemented with 2 µM Thiazovivin for 24 h. Directed cardiac differentiation was performed using a Wnt-signaling modulation protocol **(Methods S2-S5 and Figure 1B)** (*18*).

**Figure 1:**
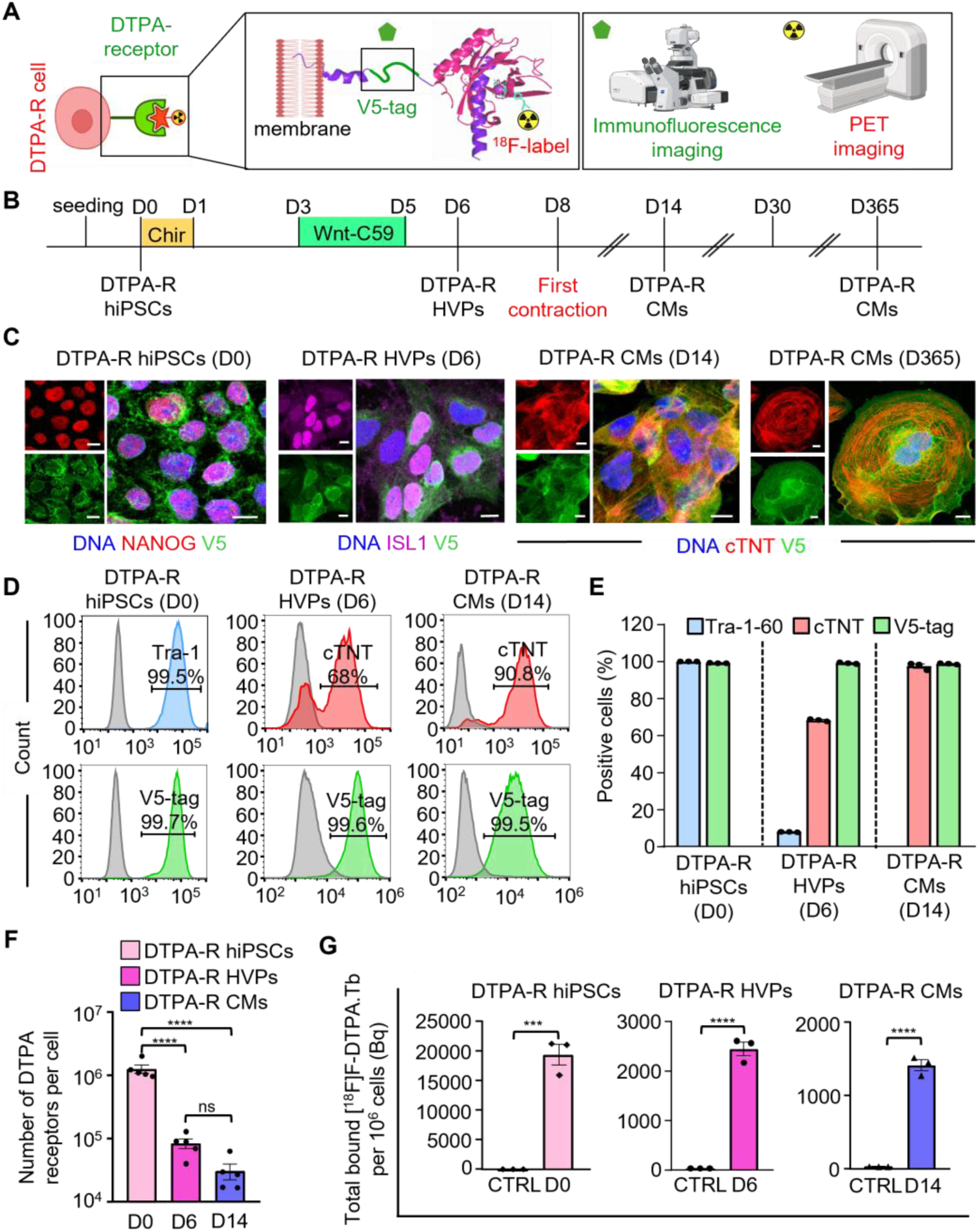
Characterization of the DTPA-R hiPSC line and its cardiac lineage. **A,** Schematic representation of the DTPA-R reporter architecture and complementary imaging readouts. The membrane-anchored DTPA-R contains an extracellular anticalin domain that binds the radiotracer [¹⁸F]F-DTPA•Tb for PET imaging and a V5-tag for immunofluorescence detection. **B,** Timeline of cardiac differentiation of DTPA-R hiPSCs into HVPs (day 6) and CMs, with onset of spontaneous contraction at day 8. **C,** Representative immunofluorescence images at the indicated differentiation stages. hiPSCs (day 0) are stained for NANOG (red), HVPs (day 6) for ISL1 (magenta), and CMs (days 14 and 365) for cTNT (red). V5-tag staining is shown at all stages (green). Nuclei are labeled with Hoechst 33258 (blue). Scale bars, 10 µm; n=3. **D,** Representative flow cytometry histograms of TRA-1-60 in hiPSCs (day 0) and HVPs (day 6), cTNT in HVPs (day 6) and CMs (day 14), with V5-tag shown at all stages. Gray histograms indicate IgG controls. **E,** Quantification of marker-positive cells from (D) (n=3). **F,** Quantification of DTPA-R receptor numbers per cell across differentiation by flow cytometry (n=5). **G,** [^18^F]F-DTPA•Tb total binding per 1×10^6^ cells at indicated stages (n=3).

### Immunofluorescence and flow cytometry analysis

Immunofluorescence analysis was performed in 2D and 3D cultures to confirm continued DTPA-R expression across differentiation stages, using the pluripotency markers OCT4 and NANOG for hiPSCs, the early cardiac markers ISL1, NKX2-5, and cTNT for HVPs, and cTNT and α-actinin to confirm CM identity and sarcomeric organization **(Method S6, Table S1)**.

Flow cytometry analysis was performed to assess expression of the pluripotency marker TRA-1-60 in DTPA-R hiPSCs and HVPs (day 6), while cTNT was analyzed in HVPs (day 6) and CMs (day 14) **(Method S7, Table S1)**. DTPA-R surface expression was quantified at all differentiation stages through detection of the V5-tag using MESF bead calibration for flow cytometry **(Method S8)** (*19*).

### *In vitro* cell labeling assays with [¹⁸F]F-DTPA•Tb

The synthesis and radiofluorination of the PET ligand for DTPA-R, [¹⁸F]F-DTPA•Tb, were performed according to established procedures reported previously (*15*) **(Method S9)**. Live DTPA-R hiPSCs, HVPs, and CMs were prepared as single-cell suspensions and incubated for 20-30 min at 37°C with [^18^F]F-DTPA•Tb in PBS + 2% BSA at a defined activity of 1 MBq per 1×10^6^ cells in a total volume of 1 mL. Following incubation, cells were washed, and the cell-bound activity was measured in a gamma counter (Wizard², PerkinElmer) **(Method S10)**.

### Effect of papain dissociation on DTPA-R surface expression

Papain is commonly used to dissociate cultivated cardiac cells prior to *in vivo* transplantation. However, proteases may potentially cleave DTPA-R at the cell surface (*19*). We therefore analyzed the effect of papain on surface DTPA-R expression by immunofluorescence and flow cytometry. Following dissociation with papain **(Method S11)**, cells were replated onto chamber slides (Lab-Tek™ II Chamber Slide System, Thermo Fisher Scientific) or 12-well plates, fixed at 12, 24, 48, and 72 h, and stained for the V5-tag and cTNT **(Method S11)**.

### Cell migration assay

Cell migration was assessed using a transwell-based fluorometric assay (Cell Migration Assay Kit, Cell Biolabs, Inc.) according to the manufacturer’s instructions, with minor adaptations as described in Supplementary Methods **(Method S12)**.

### Calcium imaging of DTPA-R CMs in 2D culture

Calcium imaging was performed on CTRL CMs and DTPA-R CMs (day 30) cultured as 2D monolayers. Intracellular calcium transients were recorded using Fluo-8 AM during electrical field stimulation at 0.5, 1, 2, and 3 Hz. Calcium transient amplitude (ΔF/F₀) was quantified using R (R Foundation for Statistical Computing) **(Method S13)**.

### 3D culture of DTPA-R HVPs on native porcine myocardial slices

CTRL HVPs and DTPA-R HVPs were seeded onto native myocardial slices at a density of 1×10^6^ cells per slice and cultured in RPMI without insulin, supplemented with 10 µM ROCK inhibitor Y-27632 for 24 h. Cultures were then maintained in RPMI without insulin, followed by a transition to RPMI containing insulin from day 8 onward.

Cell-seeded native myocardial slices were maintained for up to 14 days under biomimetic culture conditions, and contractile force generation was recorded (*4*). At experimental endpoints, slices were fixed and immunostained **(Methods S6 and S14)**.

### Repopulation of decellularized myocardial ECM with DTPA-R HVPs

Decellularized myocardial ECM scaffolds were generated from porcine left ventricular myocardium. Cell-seeded ECM constructs were cultured for up to 14 days under biomimetic conditions using the biomimetic chamber system (*20*). At experimental endpoints, constructs were fixed and immunostained **(Methods S6 and S15)**.

### [^18^F]F-DTPA•Tb PET imaging of porcine myocardial slices

Porcine myocardial slices (non-injured) were cultured in biomimetic chambers **(Method S14)**. Then, DTPA-R HVPs were seeded onto slices in the following configurations: (i) 1×10⁶ cells distributed across three seeding sites, (ii) 2×10⁶ cells distributed across two seeding sites, and (iii) 3×10⁶ cells applied at a single site. Cells were cultured on myocardium for 3 days before PET imaging.

In a separate experiment, myocardial injury was induced in the porcine slices from two individual hearts by radiofrequency ablation (RFA) **(Method S16)** on the day of cell seeding. DTPA-R HVPs (1×10^6^ cells per slice) were applied to the region opposite the injury site.

Matching RFA-injured slices without HVPs served as controls. Separate cohorts were cultured for 2 or 5 days before PET imaging.

For PET imaging, slices were incubated for 20 min at 37°C in the biomimetic chambers with 2 mL of [18F]F-DTPA•Tb at a final activity concentration of 1 MBq mL^−1^ in labeling buffer (PBS + 2% BSA). Chambers were subsequently washed six times and imaged with an Inveon (Siemens Preclinical Solutions) small-animal PET/CT scanner **(Method S17)**.

### Effect of [¹⁸F]F-DTPA•Tb on DTPA-R-expressing cells

To evaluate the effect of [^18^F]F-DTPA•Tb on DTPA-R HVPs and CMs, cells were dissociated into single-cell suspensions and incubated with 1 MBq [^18^F]F-DTPA•Tb per 1×10^6^ cells for 30 min. After washing, cell-bound radioactivity was measured from an aliquot of cells, and the remaining cells were either replated or passed once through a 29-gauge needle to impose additional mechanical stress. Controls were processed in parallel without exposure to radioactivity. Brightfield imaging was performed at 24, 48 and 72 h to assess cell attachment, morphology, and, for DTPA-R CMs, contractile activity. Cell death was assessed by collecting non-adherent cells from the supernatant and quantifying them by Trypan blue staining.

### Phantom imaging studies with a clinical PET/MR scanner

To assess PET signal detectability under clinically relevant conditions, a PET/MR phantom experiment was performed using an anthropomorphic phantom filled with 10 liters of water, with a centrally positioned compartment corresponding to the cardiac region. Two sealed samples containing identical total radioactivity (10 kBq of [^18^F]FDG) were prepared in two volumes. A 200 µL sample represented a compact intramyocardial injection volume, whereas a 1,000 µL sample represented a distribution volume due to cell migration *in vivo*. Imaging was performed on a clinical 3-T Biograph mMR PET/MR scanner (Siemens Healthineers) **(Method S18)**.

### Statistical analysis

Data are presented as mean ± SEM. Statistical analyses were performed using GraphPad Prism (v. 9.3.1; GraphPad). For two-group comparisons, two-tailed unpaired Student’s t-tests were used; for comparisons involving more than two groups, one-way or two-way ANOVA was used.

When appropriate, ANOVA was followed by post hoc multiple comparison testing as specified. Statistical significance was defined as P<0.05 (*), P<0.01 (**), P<0.001 (***), and P<0.0001 (****). Schematic images were created using BioRender (Seyfi, S. 2026; https://BioRender.com/huj2qgz) or Sarah Luger (momentsofaha.com).

## Results

### Expression of DTPA-R is maintained in hiPSC lines during differentiation

Clonal isolation yielded monoallelic and biallelic knock-in hiPSC lines with precise on-target integration and preserved genomic integrity, as confirmed by genotyping and G-band karyotyping **(Figure S1)**. Based on differentiation efficiency, a heterozygous DTPA-R hiPSC clone was selected for further experiments.

Differentiation into HVPs (day 6) and CMs was confirmed by stage-specific marker expression and the onset of spontaneous contraction at day 8. hiPSCs expressed the pluripotency markers NANOG and OCT4; HVPs expressed the cardiac progenitor markers ISL1 and NKX2-5, with cTNT already detectable at day 6; and differentiated CMs expressed cTNT, MLC2v, and sarcomeric α-actinin **(Figures 1B-C and S2)**. Immunofluorescence and flow cytometry demonstrated persistent cell-surface expression of DTPA-R across all differentiation stages, including CMs subjected to maturation for up to 365 days (Figures 1C and S2). More than 99% of cells remained V5-tag-positive throughout differentiation **(Figure 1D-E)**.

Quantification of DTPA-R revealed a progressive decrease in receptor number per cell during cardiac differentiation **(Figure 1F)**. hiPSCs exhibited mean levels of approximately 1.2×10⁶ receptors per cell, which declined to approximately 8.3×10⁴ receptors per cell in HVPs and 3×10⁴ receptors per cell in CMs.

Cellular labeling with [¹⁸F]F-DTPA•Tb, normalized to 1×10⁶ cells, followed the same trend but showed smaller relative changes **(Figure 1G)**. hiPSCs and HVPs showed radiotracer binding levels of approximately 20,000 and 2,000 Bq per 10⁶ cells, respectively, whereas CMs exhibited approximately 1,400 Bq per 10⁶ cells. Despite the reduction during differentiation, labeling of DTPA-R-expressing cells remained more than 45-fold higher than that of control cells, indicating preserved sensitivity for PET detection.

### Stability of membrane-bound DTPA-R after enzymatic dissociation of HVPs

Flow cytometry analysis of HVPs immediately after dissociation with the thiol protease papain showed that the V5-tag signal was completely lost, whereas it remained detectable after treatment with serine protease trypsin **(Figure S3)**, suggesting specific proteolytic cleavage of the extracellular reporter domain by papain. Because papain has been shown to preserve the viability of hiPSC-derived cardiac cells during dissociation, an important consideration for *in vivo* transplantation (*21*), we examined the kinetics of receptor re-expression and cardiac differentiation. Following papain dissociation and replating, DTPA-R surface expression progressively recovered over 12-72 h, as measured by flow cytometry and immunofluorescence **(Figure 2A-D)**.

**Figure 2:**
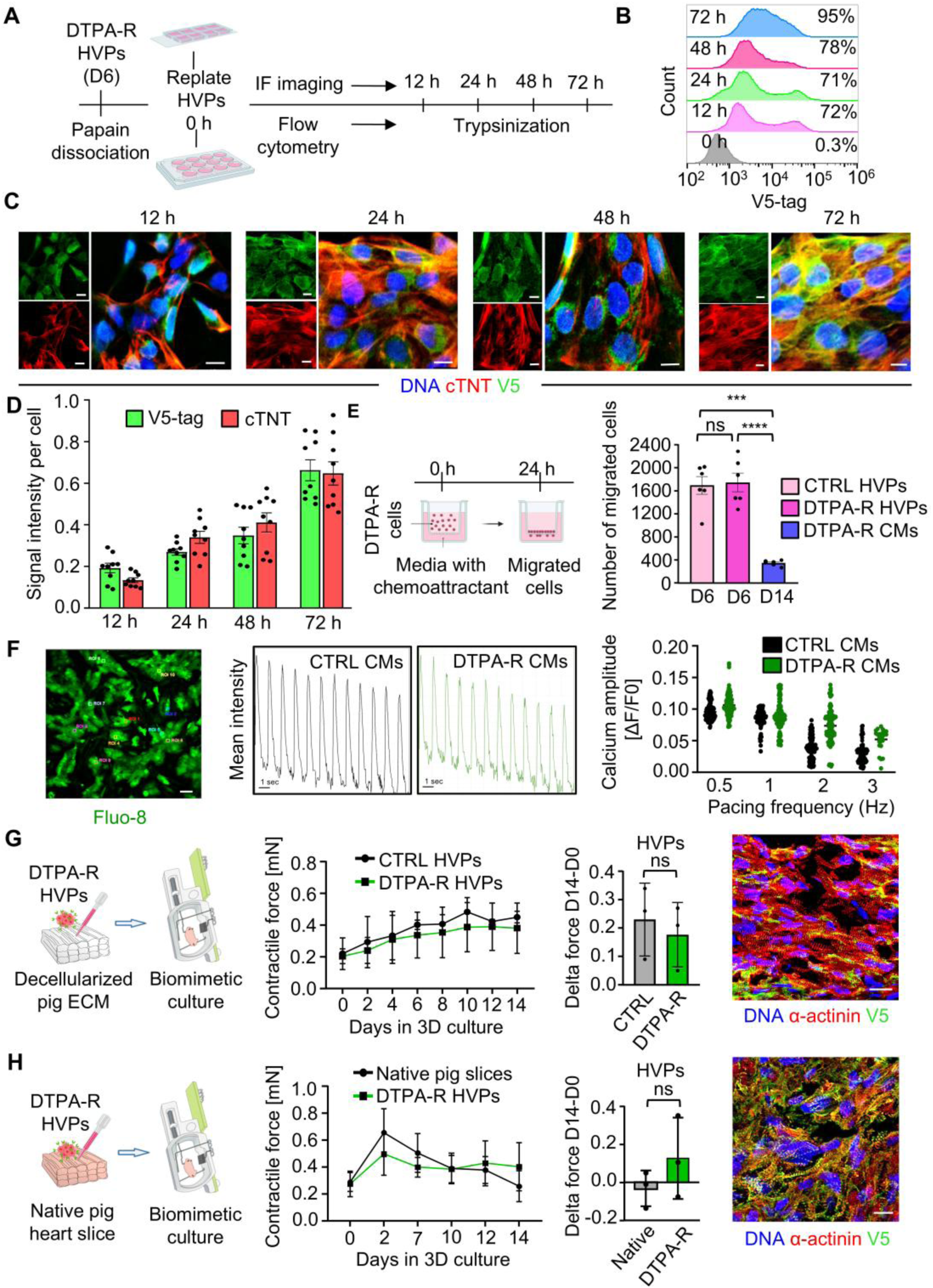
Functional characterization of DTPA-R HVPs and CMs. **A,** Experimental workflow for analysis of DTPA-R re-expression following papain dissociation. **B,** Flow cytometry histograms of surface V5-tag expression in DTPA-R HVPs at the indicated time points after replating. **C,** Representative immunofluorescence images of DTPA-R HVPs (day 6) at the indicated time points after replating, stained for V5-tag (green) and cTNT (red). Nuclei are labeled with Hoechst 33258 (blue). Scale bars, 10 µm. **D,** Quantification of V5-tag and cTNT fluorescence intensity per cell from (C) (n=3). **E,** Chemotactic migration assay. Left, schematic of the transwell setup. Right, quantification of migrated cells after 24 h. **F,** Calcium transient analysis in CMs. Left, representative image of Fluo-8–loaded CMs with regions of interest indicated (ROIs), scale bar, 100 µm. Middle, representative calcium transients of DTPA-R and CTRL CMs. Right, quantification of calcium transient amplitude (ΔF/F₀) at increasing pacing frequencies. Data represent 10 ROIs per sample (n=3). **G and H,** Functional analysis of DTPA-R HVPs (day 6) in decellularized cardiac ECM (G) and native porcine myocardial slices (H). Left, schematic of biomimetic culture; middle, contractile force over time; right, change in force between day 14 and day 0 (D14-D0) and representative immunofluorescence at day 14, with ECM patches (G) and slices (H) stained for α-actinin (red) and V5-tag (green). Scale bars, 30 µm (n=3).

### DTPA-R expression does not impair progenitor migration, differentiation, or functional integration

Migration capacity was comparable between DTPA-R and CTRL HVPs, whereas DTPA-R CMs showed markedly lower migration, consistent with the loss of progenitor migratory behavior during differentiation into cardiomyocytes **(Figure 2E)**. Calcium imaging demonstrated preserved excitation-contraction coupling in DTPA-R CMs, with the expected frequency-dependent reduction in calcium transient amplitude across increasing pacing frequencies **(Figure 2F)**.

The differentiation capacity and functional performance of DTPA-R HVPs were further evaluated in physiologically relevant 3D culture models. In decellularized porcine cardiac ECM constructs cultured under mechanical loading and electrical pacing, DTPA-R HVP–derived tissues developed increasing contractile force over 14 days, comparable to CTRL HVP-seeded constructs **(Figure 2G)**. Immunofluorescence confirmed differentiation into CMs and persistent V5-tag expression throughout the engineered cardiac tissue.

Functional integration was additionally assessed in native porcine myocardial slices. DTPA-R HVP-seeded slices maintained contractile activity beyond day 10, whereas control slices showed the expected progressive decline in contractile performance **(Figure 2H)**. Immunofluorescence further demonstrated V5-tag and sarcomeric α-actinin expression in DTPA-R cells within the myocardial tissue.

### *Ex vivo* PET imaging enables detection of DTPA-R HVPs in myocardial tissue

Slices without seeded cells showed only background PET signal across all experiments, confirming assay specificity. In non-injured myocardium, all HVP-seeded conditions produced focal PET signals at the deposition site. Already 1×10^6^ cells distributed across three sites were detectable **(Figure S4)**. In RFA-injured myocardium, PET imaging on days 2 and 5 post-injury showed a discrete focal [^18^F]F-DTPA•Tb signal specifically in slices seeded with DTPA-R HVPs **(Figure 3)**. By day 5, signal was detected not only at the original seeding site but also at the injury site, indicating that cells had migrated toward the lesion **(Figure S5)**. Post-imaging immunofluorescence confirmed these findings: V5-tag-positive cells were present within and around injury regions at both time points, despite having been seeded at a remote location, consistent with migration toward the lesion **(Figure 3F-G)**. Notably, these migrating cells co-expressed cTNT, suggesting they had undergone differentiation within the injured tissue. Control injury samples showed no V5-tag signal in immunofluorescent staining **(Figure S6)**.

**Figure 3:**
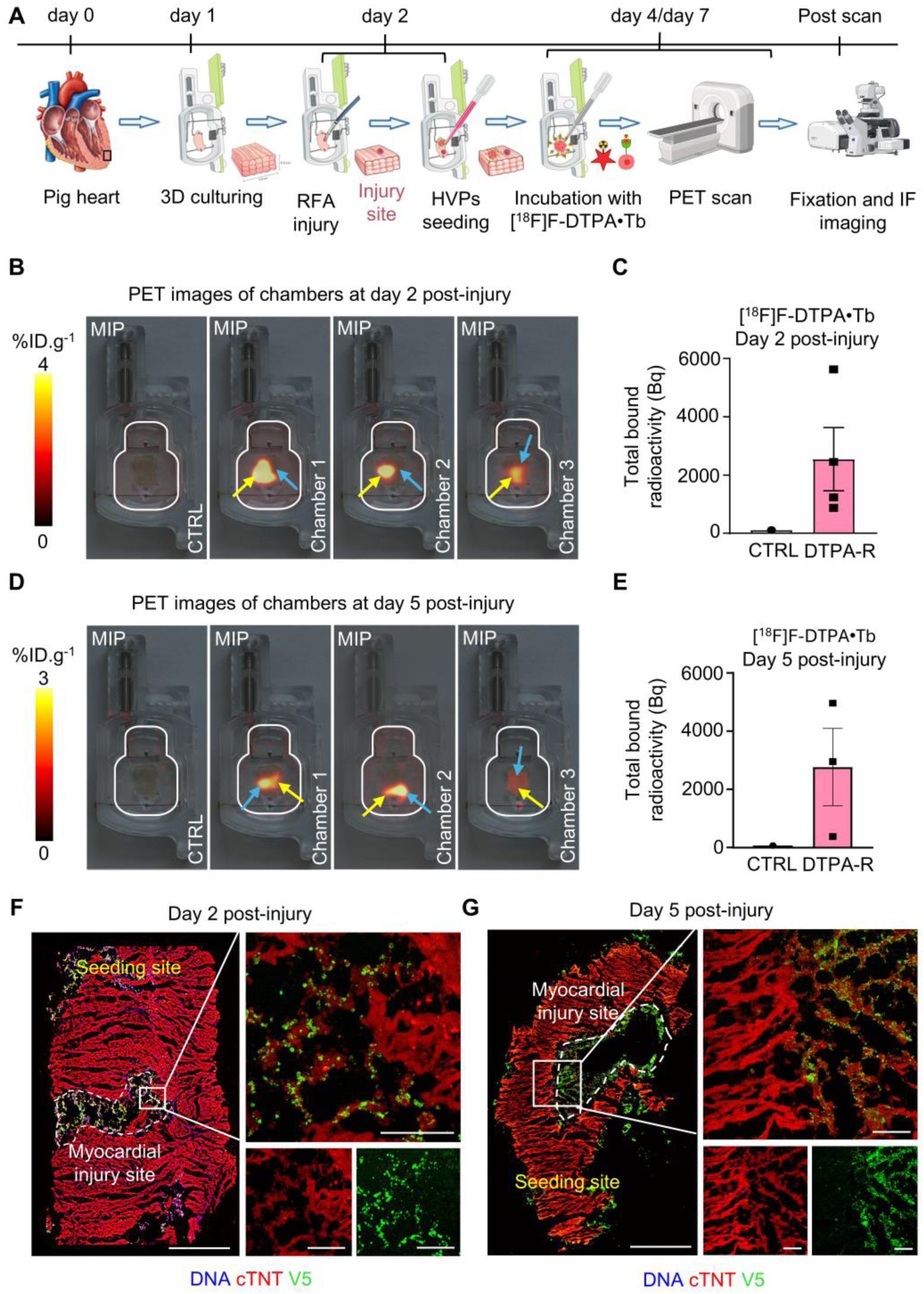
*Ex vivo* PET detection of DTPA-R HVPs in RFA injury model. **A,** Schematic of the *ex vivo* RFA-injury model and imaging workflow. The RFA-injury model was established using native porcine myocardial slices from two individual hearts. Myocardial slices from one heart were imaged on day 2 post-injury, and slices from the second heart were imaged on day 5 post-injury. **B and D,** Maximum-intensity-projection (MIP) PET images of RFA-injured tissue chambers on day 2 (B) and day 5 (D) post-injury, including a control chamber without cells (CTRL) and chambers seeded with DTPA-R HVPs (Chambers 1–3). Yellow arrows indicate the cell seeding sites, and blue arrows indicate the corresponding myocardial injury sites. **C and E,** Total PET-detected radioactivity (Bq) per chamber on day 2 (C) (CTRL, n=1; DTPA-R, n=4) and day 5 (E) (CTRL, n=1; DTPA-R, n=3) post-injury. **F and G,** Representative immunofluorescence tile-scan images of RFA-injured tissue on day 2 and day 5 post-injury after PET imaging, stained for cTNT (red) and V5-tag (green). Nuclei are labeled with Hoechst 33258 (blue). Left, tile-scan (merged); right, higher-magnification views of boxed regions. Scale bars, 1 mm (overview), 100 µm (magnified).

### Exposure to [^18^F]F-DTPA•Tb does not impair cell viability or phenotype

For *in vivo* transplantation, cardiac cells are exposed to the radiotracer for imaging and delivered intramyocardially through a syringe needle. We therefore examined the effects of [¹⁸F]F-DTPA•Tb exposure and syringe-based handling on the viability, morphology, and phenotype of freshly thawed HVPs and CMs over 72 h **(Figure 4A)**. Gamma-counter measurements confirmed [¹⁸F]F-DTPA•Tb binding to DTPA-R-expressing cells **(Figure 4B)**. Bright-field imaging at 24 h showed increased detachment in DTPA-R CMs following syringe passage compared to HVPs, whereas DTPA-R HVP morphology remained largely preserved **(Figure 4C)**. Cell death measurement revealed a transient increase in the dead-cell fraction at 24 h, particularly after syringe passage, followed by recovery toward baseline levels by 72 h **(Figure 4D-E)**. DTPA-R CMs showed a more pronounced response to syringe handling than DTPA-R HVPs, whereas labeled cells followed a similar overall viability pattern to the corresponding unlabeled controls. Immunofluorescence at 72 h confirmed preservation of cTNT and V5-tag expression in radiolabeled HVPs and CMs, both with and without syringe passage **(Figure 4F-G)**. Together, these findings indicate that syringe passage induced a transient reduction in cell viability, particularly in CMs, whereas [¹⁸F]F-DTPA•Tb exposure produced no obvious additional adverse effect on viability or cardiac phenotype under the conditions examined.

**Figure 4:**
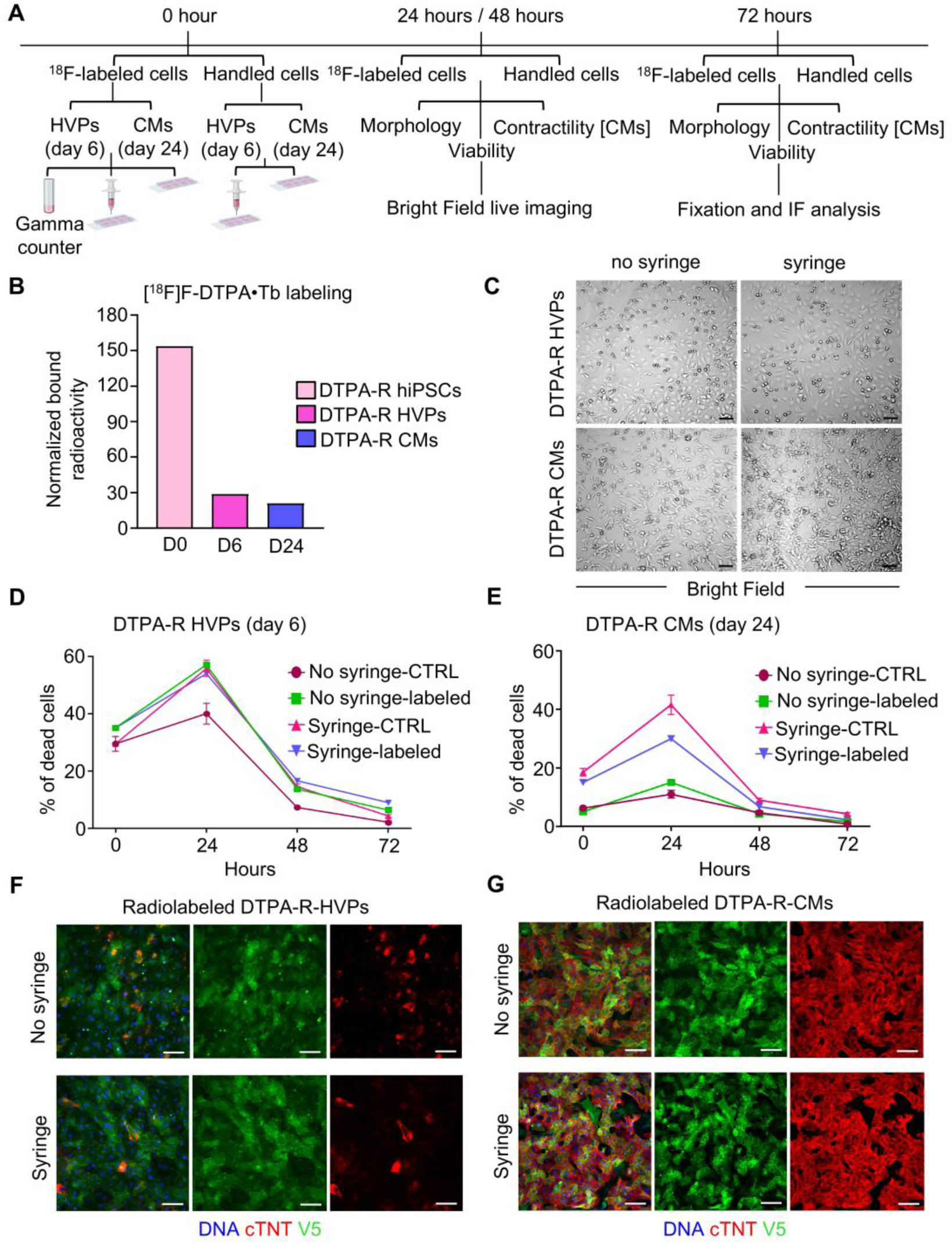
Evaluation of radiotracer exposure and stress responses in DTPA-R hiPSC-derived cardiac cells. **A,** Schematic of the experimental workflow. DTPA-R HVPs (day 6) and CMs (day 24) were exposed to [^18^F]F-DTPA•Tb or left unlabeled (CTRL), subjected to syringe passage or not, and returned to culture for 72 h. **B,** Cell-associated [^18^F]F-DTPA•Tb radioactivity expressed as fold change relative to the control (CTRL=1), measured by gamma counter for the indicated cell types (n=1). **C,** Representative bright-field images acquired 24 h after replating, showing the morphology of labeled DTPA-R HVPs and CMs handled with or without syringe passage. Scale bars, 50 µm. **D and E,** Percentage of dead DTPA-R HVPs (D) and CMs (E) measured over 72 h under the indicated conditions. For both cell types, labeled cells with or without syringe passage were analyzed (n=1 per group), whereas unlabeled control cells with or without syringe passage were analyzed (n=4 per group). **F and G,** Representative immunofluorescence images of labeled DTPA-R HVPs (F) and CMs (G) stained for cTNT (red) and V5-tag (green) at 72 h. Nuclei are labeled with Hoechst 33258 (blue). Scale bars, 100 µm.

### PET/MR phantom study

To mimic clinically relevant intramyocardial cell transplantation, a phantom study was performed to evaluate the effect of spatial distribution volume on PET signal using identical activities distributed over two injection volumes. At the first scan (0 h), ROI-based PET quantification yielded a total activity of approximately 7.8 kBq. Based on the measured mean *in vitro* cell-associated radiotracer binding values, this activity corresponded to approximately 3.2×10^6^ DTPA-R HVPs (2.4 kBq/10^6^ cells) or 5.5×10^6^ DTPA-R CMs (1.4 kBq/10⁶ cells) **(Figure 5C-D)**. These values represent the cell-equivalent numbers corresponding to the activity measured at the beginning of imaging rather than a minimum detection threshold. When displayed using identical intensity scales, the 200-µL condition consistently appeared as a compact high-contrast hotspot, whereas the 1,000-µL condition showed lower signal intensity because the same activity was distributed over a larger volume **(Figure 5A-B)**. The compact 200-µL source remained visually distinguishable at all time points, whereas the 1,000-µL source approached background signal levels by 6 h.

**Figure 5:**
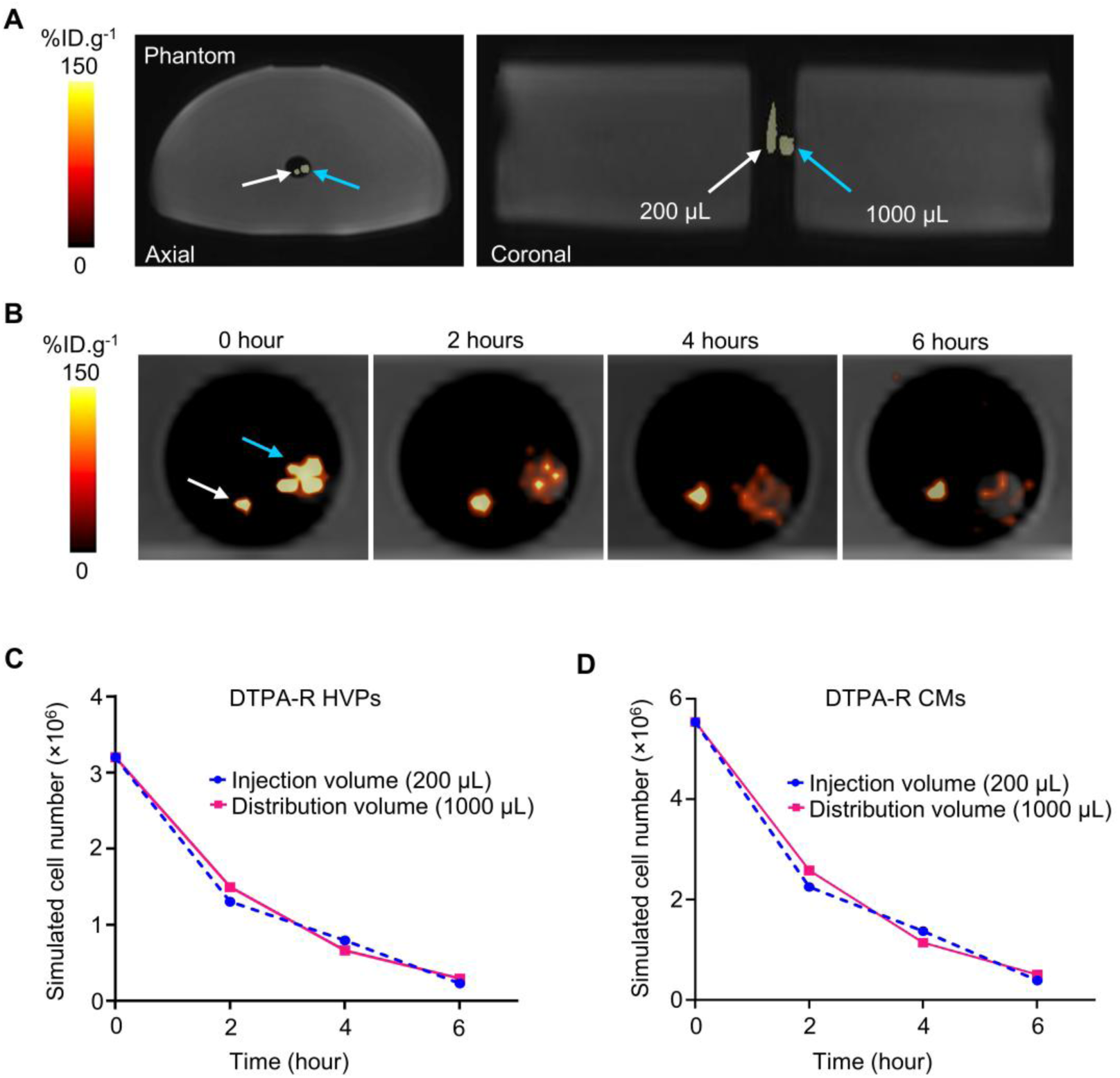
Phantom PET/MR evaluation of simulated intramyocardial cell injection volumes. **A,** PET/MR images of an anthropomorphic water-filled phantom with thoracic attenuation properties. Two sealed sources containing 10 kBq of [¹⁸F]FDG each were positioned in the central region of the phantom, corresponding to the cardiac region. One source was confined to a 200-µL injection volume (white arrow), and the other to a 1,000-µL injection volume (blue arrow). Axial and coronal PET/MR views are shown. **B,** PET images acquired at 0, 2, 4, and 6 h (20-min static acquisitions), displayed using the same intensity scale (%ID·g^−1^) for both sources. **C and D,** Total PET activity was calculated from ROIs encompassing the entire signal at each scan time point. The measured activity was converted to estimated cell-equivalent numbers of DTPA-R HVPs (C) and CMs (D) using the mean *in vitro* cell-associated [¹⁸F]F-DTPA•Tb activity (Bq/10⁶ cells) for the 200- and 1,000-µL volumes.

## Discussion

Our data indicate that expression of the reporter gene DTPA-R does not affect fundamental biological properties of hiPSCs or their derived progeny. Across 2D, 3D, and *ex vivo* models, engineered cells retained differentiation potential, migration capacity, and functional integration. This is notable because genetic manipulation has the potential to alter cellular phenotype and function (*22,23*).

Validation in physiologically relevant models is crucial. Decellularized ECM constructs enabled assessment of force generation and electromechanical coupling under defined mechanical loading, while viable myocardial slices allowed cellular integration and injury responses (*24–27*). DTPA-R HVPs demonstrated migration toward injury sites, consistent with previously reported regenerative behavior (*4*). PET imaging detected these cells *ex vivo*, highlighting the potential of this system for tracking spatial cell dynamics. Exposure to [^18^F]F-DTPA•Tb did not induce cytotoxicity, supporting the biological inertness of the DTPA-R system and reinforcing its translational suitability. HVPs tolerated radiolabeling and syringe passage substantially better than CMs, and together with their reported regenerative and engraftment potential, these results further support progenitor-based strategies for cardiac cell therapy (*4*).

Beyond biological validation, phantom experiments addressed a key translational imaging constraint by demonstrating that PET signal visibility is governed by activity per volume (concentration) rather than by the total activity. By modeling clinically relevant intramyocardial injection volumes and post-injection dispersion under large-animal or human attenuation conditions, we showed that compact cell deposits yield substantially higher apparent PET signal than distributed activity with identical tracer content in total. While fundamental to PET physics, this principle is rarely addressed explicitly in the context of cell therapy. These findings provide practical guidance for interpreting PET signal evolution as transplanted cells disperse or migrate within the tissue. The phantom experiment was performed under homogeneous background conditions. *In vivo*, inhomogeneous background from neighboring structures and cardiac motion may further reduce signal contrast and increase the number of labeled cells required for reliable PET detection.

Several limitations should be acknowledged. *In vivo* PET imaging was not performed, and thus the influence of full perfusion, immune interactions, and tracer pharmacokinetics remains to be determined. Notably, Anticalin proteins have exhibited favorable safety profiles in clinical studies after intravenous administration, including low immunogenicity (*15,28*). Nevertheless, the long-term safety of cell surface-expressed DTPA-R still has to be determined.

In conclusion, our study lays the foundation for future *in vivo* applications of a promising new reporter gene system for monitoring cardiac regenerative therapies. The system is bioorthogonal, shows no measurable impact on cardiac progenitor cell differentiation and function, is stably expressed over months, and demonstrates high specific labeling in cell culture and *ex vivo* studies.

## Acknowledgements

The authors thank Birgit Campbell and Marco Crovella for their excellent technical assistance in cell culture, Sybille Reder and Romina Karampour for PET imaging support, and Sophie Zengerle for her assistance with immunofluorescence image quantification. This work was funded by the European Research Council (grants 788381 and 101141820 to A.M.), the German Research Foundation (Transregio Research Units 152 and 267 to A.M. and K.-L.L.), the Jung Foundation for Science and Research (C.M.P.), and the German Center for Cardiovascular Research (DZHK; grant FKZ 81Z0600601 to A.M. and K.-L.L.).

## Author contributions

S.S. conceived and led the study, designed and carried out the experiments, analyzed and interpreted the data, prepared the figures, and drafted the manuscript. T.D. generated CRISPR– Cas9–engineered iPSC lines and contributed to experimental design. C.M.P. contributed to *ex vivo* heart slice experiments and experimental design. M.G. conducted PET/MR acquisitions. E.B. contributed to calcium imaging and its analysis. V.F. assisted *ex vivo* heart slice experiments. A.S. contributed to reporter design. K.F. and V.M. constructed plasmids and supported experimental strategy.

M.S., K.-L.L., C.K., A.M., and W.A.W. supervised the study. T.D., C.M.P., W.A.W., and A.M. edited the manuscript. M.S., A.M., K.-L.L., C.M.P. and W.A.W. secured funding. All authors reviewed and approved the final manuscript.

## Competing Interests

The authors declare no competing interests.

## Ethics declarations

No human participants or live animal experiments were involved. hiPSCs were used exclusively *in vitro*. Porcine hearts were obtained post mortem from animals euthanized for other approved studies and used only *ex vivo*. According to the German Animal Welfare Act (Tierschutzgesetz) and Bavarian regulations, post-mortem animal tissue does not require additional ethical approval.

## Data Availability

The data supporting the findings of this study are available from the corresponding authors, W.A.W. and A.M., upon request.

## Notes

### Competing Interest Statement

The authors have declared no competing interest.

