## Supplementary Material for "A new bioorthogonal PET reporter gene for cardiac regenerative medicine binding to DTPA-lanthanide complexes"

### Supplementary methods

#### Method S1: Generation of DTPA-R hiPSC reporter line

hiPSCs were co-nucleofected using a 4D-Nucleofector system (Lonza) with a Cas9/AAVS1 single-guide RNA plasmid (Addgene #80494) and the DTPA-R donor plasmid containing homology arms and a puromycin resistance cassette (*1*). Cells were plated on Matrigel-coated plates in mTeSR1 medium supplemented with 10 μM Y-27632 for 24 h. Puromycin selection (0.2 μg $mL^{-1}$) was initiated 48 h post-nucleofection and maintained for 5 days. Resistant colonies were manually picked, expanded, and screened for correct AAVS1 integration by junction PCR and Sanger sequencing. Predicted off-target sites were assessed by targeted PCR and sequencing (*1*). Karyotyping was performed, and a clone with no chromosomal abnormality, normal morphology, and stable growth was expanded and used for all subsequent experiments.

#### Method S2: DTPA-R hiPSC maintenance and cardiac differentiation

hiPSCs were seeded onto Geltrex-coated 24-well plates in Essential 8 (E8) medium supplemented with 2 μM Thiazovivin for 24 h. Cardiac differentiation was initiated upon reaching >95% confluency by transient Wnt activation with 1 μM CHIR99021, followed by Wnt inhibition with 2 μM Wnt-C59 at day 3 (*2*). Cells were maintained in RPMI 1640 supplemented with B27 without insulin, with medium changes every other day. HVPs were obtained at day 6 and either purified by MACS depletion of TRA-1-60-positive cells **(Method S3)** or further differentiated into CMs. From day 8 onward, CMs were cultured in RPMI 1640 supplemented with B27 containing insulin. DTPA-R HVPs and CMs were dissociated using trypsin or papain-based methods (*3*), depending on experimental requirements **(Method S4)**. Long-term maintenance of CMs for up to one year is described in **Method S5**. HVPs were cryopreserved in CryoStor CS10. All cultures were routinely confirmed to be mycoplasma free.

#### Method S3: Purification of DTPA-R HVPs by TRA-1-60 magnetic separation

To deplete residual undifferentiated pluripotent stem cells, day-6 differentiated cultures were purified by magnetic-activated cell sorting (MACS) using Anti-TRA-1-60 MicroBeads, human (Miltenyi Biotec; Cat. No. 130-100-832), adapted from the previously described HVP purification procedure and the manufacturer's instructions (*4*).

Cells were dissociated into single-cell suspensions using 0.05% trypsin and washed once with cold separation buffer consisting of Dulbecco's phosphate-buffered saline (DPBS) supplemented with 0.5% bovine serum albumin (BSA) and 2 mM EDTA. After cell counting, suspensions were centrifuged at 300×g for 5 min and resuspended in separation buffer at 80 μL per $2\times10^6$ cells. Anti-TRA-1-60 MicroBeads were added at 20 μL per $2\times10^6$ cells, followed by incubation for 5 min at 2-4 °C. The suspension was then adjusted to a final volume of 1 mL with separation buffer.

Magnetic separation was performed manually using MS Columns (Miltenyi Biotec; Cat. No. 130-042-201) placed in a MACS Separator. Columns were pre-equilibrated with 0.5 mL separation buffer before application of the labeled cell suspension. The flow-through fraction, containing TRA-1-60-negative HVPs, was collected. Columns were then washed three times with 0.5 mL separation buffer, and all flow-through fractions were pooled. TRA-1-60-positive cells were retained in the column and discarded.

To assess the efficiency of TRA-1-60 depletion, flow cytometry was performed on purified cell fractions in representative experiments to determine the proportion of residual TRA-1-60-positive cells. Routine flow-cytometric validation was not performed for every preparation.

Following separation, collected TRA-1-60-negative cells were centrifuged, washed once with DPBS, and immediately cryopreserved to preserve the day-6 progenitor state.

**Method S4: DTPA-R HVPs and CMs dissociation methods**

For enzymatic dissociation with trypsin, HVPs were incubated with 0.05% trypsin (Gibco), whereas CMs were incubated with 0.25% trypsin (Gibco). In both cases, approximately 0.5 mL trypsin solution per well of a 24-well plate was applied, and cells were incubated at 37°C for 8-12 min. Enzymatic activity was terminated by the addition of trypsin inhibitor (Sigma-Aldrich; T9253) at a final concentration of 0.5-1 mg $mL^{-1}$, followed by gentle trituration to obtain a single-cell suspension.

For experiments requiring enhanced preservation of cell viability, papain-based dissociation was employed as previously described (*3*). Cells were washed twice with 2 mM EDTA in calcium- and magnesium-free PBS. A 2× papain solution was freshly prepared with papain suspension (Worthington Biochemical Corporation; LS003124) at 40 U $mL^{-1}$ and 2 mM L-cysteine hydrochloride monohydrate (Sigma-Aldrich; C6852) in calcium- and magnesium-free

PBS and incubated at 37°C for 10 min. The 2× papain solution was then diluted 1:2 in PBS to obtain the 1× papain solution before use. Cells were incubated at 37°C for 20 min for 2D cultures or 25-30 min for spheroids, with optional shaking. Digestion was terminated by the addition of an equal volume of stop solution containing 1 mg $mL^{-1}$ trypsin inhibitor (Sigma-Aldrich; T9253) and 40 µg $mL^{-1}$ DNase I (Sigma-Aldrich; DN25). Cells were gently dissociated by pipetting, washed with culture medium, centrifuged at 250×g for 5 min, and resuspended in the desired culture medium.

**Method S5: Long-term maintenance of DTPA-R CMs**

For long-term CM culture (typically ⩾2 months after differentiation), cultures were examined by light microscopy to identify regions exhibiting stable, synchronous contractions. Beating regions were mechanically excised by cutting around the contractile area using a sterile insulin needle. Excised cardiomyocyte clusters were gently aspirated using a 1 mL pipette tip, transferred into microcentrifuge tubes, and allowed to settle by gravity. After removal of the supernatant, clusters were replated onto fibronectin-coated 12-well plates.

For the first 24 h after replating, cells were maintained in EB20 medium, consisting of DMEM/F-12 supplemented with 20% fetal bovine serum (FBS), 1% L-glutamine, 1% non-essential amino acids, 0.5% penicillin/streptomycin, and 0.1 mM β-mercaptoethanol. After the initial recovery period, the medium was replaced with EB2, consisting of DMEM/F-12 supplemented with 2% FBS, 1% L-glutamine, 1% non-essential amino acids, 0.5% penicillin/streptomycin, and 0.1 mM β-mercaptoethanol. CMs were subsequently maintained in EB2 with partial medium changes every few days and were cultured for up to one year.

**Method S6: Immunofluorescence analysis of 2D and 3D cultures**

Immunofluorescence staining was performed on 2D cell cultures and 3D tissue samples to assess pluripotency, cardiac lineage commitment, structural maturation, and DTPA-R reporter expression. A complete list of primary and secondary antibodies, including antigen, clone, vendor, and dilution, is provided **(Supplementary Table S1)**.

For 2D staining, hiPSCs were seeded directly onto Geltrex™-coated chamber slides (Geltrex LDEV-Free; Thermo Fisher Scientific). HVPs and CMs were dissociated into single cells using either 0.05% or 0.25% trypsin or papain, as described above, and subsequently replated onto chamber slides prior to staining. HVPs were plated onto Geltrex™-coated chamber slides, whereas CMs were plated onto fibronectin-coated chamber slides. Following papain-based dissociation, CMs were allowed to recover for 3-4 days before immunofluorescence analysis. Cells were fixed with 4% paraformaldehyde (PFA; Sigma-Aldrich) prepared in Dulbecco's phosphate-buffered saline containing calcium and magnesium (DPBS with $Ca^{2+}/Mg^{2+}$; Gibco) for 10-15 min at room temperature and washed thoroughly with the same buffer. Permeabilization was performed for 10 min using DPBS with $Ca^{2+}/Mg^{2+}$ supplemented with 0.1% Triton X-100 (Sigma-Aldrich). Samples were blocked for 1 h at room temperature in DPBS with $Ca^{2+}/Mg^{2+}$ containing 10% fetal bovine serum (FBS). Primary antibodies were diluted in DPBS with $Ca^{2+}/Mg^{2+}$ containing 0.5% bovine serum albumin (BSA) and incubated overnight at 4°C. After washing with DPBS with $Ca^{2+}/Mg^{2+}$ containing 0.1% Triton X-100, fluorophore-conjugated secondary antibodies were diluted in DPBS with $Ca^{2+}/Mg^{2+}$ containing 0.5% BSA and incubated for 1 h at room temperature. Final washes were performed using DPBS with $Ca^{2+}/Mg^{2+}$ without detergent. Nuclei were counterstained with Hoechst 33258 at 5 $\mu g\ mL^{-1}$, and samples were mounted using ProLong™ Gold Antifade Mountant.

For 3D tissue immunofluorescence, samples were fixed in 4% PFA prepared in DPBS with $Ca^{2+}/Mg^{2+}$ for 24-48 h at 4°C, with the fixation period adjusted according to sample size. Fixed samples were cryoprotected in 20% sucrose, embedded in Tissue-Tek O.C.T. compound (Sakura Finetek), frozen in 2-methylbutane chilled with liquid nitrogen, and cryosectioned at 12 μm thickness. Tissue sections were permeabilized using DPBS with $Ca^{2+}/Mg^{2+}$ supplemented with 0.3% Triton X-100 and blocked in DPBS with $Ca^{2+}/Mg^{2+}$ containing 10% FBS. Primary antibodies were diluted in DPBS with $Ca^{2+}/Mg^{2+}$ containing 0.5% BSA and incubated overnight at 4°C. After washing with DPBS with $Ca^{2+}/Mg^{2+}$ containing 0.1% Triton X-100, fluorophore-conjugated secondary antibodies were diluted in DPBS with $Ca^{2+}/Mg^{2+}$ containing 0.5% BSA and incubated for 1 h at room temperature. Nuclei were counterstained with Hoechst 33258 at 5 $\mu g\ mL^{-1}$, and sections were mounted using ProLong™ Gold Antifade Mountant.

Fluorescence imaging was performed using a Thunder imaging system or a Leica SP8 confocal microscope (Leica Microsystems). Image acquisition was conducted using LAS X software (Leica Microsystems), and image processing and quantitative analyses were performed using Fiji/ImageJ. Quantification included assessment of marker-specific fluorescence signal intensity on a per-cell basis.

**Method S7: Flow cytometry**

For flow cytometric analysis, single-cell suspensions were prepared in FACS buffer (PBS supplemented with 1% fetal bovine serum [FBS]). Cells were fixed with 4% paraformaldehyde (PFA) in PBS for 10-15 min at room temperature. Surface staining was performed using anti-TRA-1-60 and anti-V5-tag antibodies on live or fixed, non-permeabilized cells, as appropriate. Cells were incubated with the respective antibodies in FACS buffer for 40-60 min at 4°C, washed three times with FACS buffer, and resuspended in FACS buffer for acquisition. For intracellular cTNT staining, fixed cells were permeabilized and blocked in PBS containing 10% FBS and 0.1% Triton X-100 before incubation with anti-cTNT antibody in PBS containing 1% FBS and 0.1% Triton X-100. Corresponding IgG isotype controls were included for all stainings. Antibody details are provided (Supplementary Table S1).

Prior to acquisition, all samples were washed and passed through a 40-μm cell strainer to remove aggregates. Data were acquired on a Gallios flow cytometer (Beckman Coulter) and analyzed using FlowJo software (v10.8.1; Becton Dickinson). Marker expression was quantified as the percentage of positive cells and/or mean fluorescence intensity, as indicated.

**Method S8: Assessment of DTPA-R expression via the V5-tag**

Single-cell suspensions of DTPA-R hiPSCs, HVPs, and CMs were prepared and stained as described under Flow Cytometry. V5-tag expression was detected using an Alexa Fluor 488–conjugated anti-V5-tag antibody, and median fluorescence intensity (MFI) was determined from gated singlet populations after subtraction of background fluorescence from V5-tag-negative controls (*5*). Calibration was performed using Quantum™ MESF Alexa Fluor 488 beads (Bangs Laboratories) acquired under identical instrument settings. Interpolated MESF values were used to calculate receptor numbers per cell according to the following equation:

$$Receptors\ per\ cell\ = \frac{\left(MESF_{sample} - MESF_{control}\right)}{(DOL)} \times 2$$

where MESF_sample and MESF_control represent interpolated MESF values of V5-tag-positive and V5-tag-negative cells, respectively, and DOL denotes the experimentally determined antibody degree of labeling. Data were analyzed using FlowJo software.

**Method S9: Chemical synthesis of precursors and radiofluorination of [$^{18}$F]F-DTPA•Tb**

Briefly, the non-radioactive precursor TMA-Nic-d-Glu$_2$-PEG$_4$-CHX-A″-DTPA was synthesized using combined solid- and solution-phase chemistry, conjugated to a DTPA chelator, and purified and characterized as described in the original report (*6*).

[$^{18}$F]Fluoride was produced via the $^{18}$O(p,n)$^{18}$F reaction using a PETtrace 880 cyclotron (GE Healthcare) and radiolabeled on a Modular-Lab Standard synthesis module (Eckert & Ziegler). Following radiolabeling, the product was purified by reversed-phase HPLC, complexed with terbium(III), and formulated in DPBS, yielding [$^{18}$F]F-DTPA•Tb for immediate use in downstream experiments.

**Method S10: Measurement of cell-bound [$^{18}$F]F-DTPA•Tb**

DTPA-R hiPSCs, HVPs, and CMs were prepared as single-cell suspensions and resuspended in PBS containing 2% BSA. Cells were incubated with [$^{18}$F]F-DTPA•Tb at an activity of 1 MBq per $1\times10^6$ cells in a final volume of 1 mL for 20-30 min at 37°C.

Cells were washed three times with PBS containing 2% BSA to remove unbound radiotracer and recounted to determine viable cell numbers. Cell-associated radioactivity was measured using a gamma counter (Wizard$^2$, PerkinElmer). Counts were converted to absolute activity values using a standardized reference measurement acquired prior to the experimental series and applied consistently across samples. Radiotracer uptake was reported as total bound radioactivity or as fold change relative to control samples.

**Method S11: V5-tag receptor re-expression assay**

Time-dependent re-expression of surface DTPA-R following enzymatic cleavage was assessed in DTPA-R HVPs (day 6) using immunofluorescence and flow cytometry. HVPs were dissociated using papain, which cleaves the extracellular domain of DTPA-R, resulting in loss of detectable surface V5-tag (*5*), and replated under standard progenitor culture conditions.

For immunofluorescence analysis, cells were replated onto Geltrex™-coated chamber slides (Lab-Tek™ II Chamber Slide System; Thermo Fisher Scientific), fixed at 12, 24, 48, and 72 h, and stained for V5-tag and cTNT. Nuclei were counterstained with Hoechst 33258. Receptor re-expression kinetics were quantified based on V5-tag fluorescence intensity per cell, while cTNT signal intensity per cell was used to monitor the progression of cardiac differentiation.

In parallel, cells replated in Geltrex™-coated 12-well plates were dissociated at matching time points using 0.05% trypsin to generate single-cell suspensions for flow cytometry. Surface V5-tag expression was quantified as the percentage of V5-tag-positive cells, and data were analyzed using FlowJo software.

**Method S12: Cell migration assays**

CTRL HVPs (d6) and DTPA-R HVPs (d6) were dissociated with trypsin, and DTPA-R CMs (d14) were dissociated with papain, resuspended in serum-free medium, and seeded ($0.5\times10^6$ cells) into transwell inserts with 8-μm pore size membranes. The lower chamber contained basal RPMI medium supplemented with SDF-1 (80 ng $mL^{-1}$). After 24 h incubation at 37°C and 5% $CO_2$, non-migrated cells were removed, and migrated cells were processed using the kit's fluorometric detection reagents. Fluorescence images were acquired using a Thunder microscope, and migrated cells were quantified by counting fluorescence-positive cells using Fiji/ImageJ.

**Method S13: Calcium imaging of DTPA-R CMs in 2D culture**

Calcium imaging was performed on CTRL CMs and DTPA-R CMs (day 30) cultured as 2D monolayers. Cells were dissociated using papain and seeded onto fibronectin-coated glass coverslips (~$9\times10^4$ cells per coverslip) and maintained in EB2 medium until the formation of a confluent, spontaneously beating monolayer. Coverslips were transferred to a temperature-controlled imaging chamber and maintained at 37°C in EB2 medium prior to imaging. Cells were loaded with 4 μM Fluo-8 AM in EB2 medium for 30 min at 37°C, washed, and equilibrated in 2 mL Tyrode's buffer solution, which was used for all imaging experiments. Intracellular calcium transients were recorded using Fluo-8 AM (AAT Bioquest) on a Thunder microscope during electrical field stimulation at 0.5, 1, 2, and 3 Hz. Fluorescence time-series data were analyzed using R (R Foundation for Statistical Computing). Calcium transient amplitude was quantified as $\Delta F/F_0$.

**Method S14: Biomimetic 3D culture of porcine myocardial slices**

Porcine ventricular myocardial slices were generated and maintained under biomimetic culture conditions, following the established methodology described previously (*4*). Fresh porcine ventricular myocardium was excised and prepared into tissue blocks, from which myocardial slices with dimensions of approximately 1.0 cm × 0.5 cm and a thickness of ~300 μm were generated, using a vibratome (Leica). For *ex vivo* culture, myocardial slices were cultured in MyoDish biomimetic culture chambers (InVitroSys, Germany). Both ends of each slice were mechanically anchored to polyethylene terephthalate support elements using tissue adhesive (Histoacryl, B. Braun), enabling defined physiologic preload. Cultured slices were assembled within the chamber configuration as previously described. Chambers were filled with a myocardial slice culture medium M199 supplemented with ITS suitable for large-mammal cardiac tissue, prepared and used according to established protocols. Cultures were maintained in a humidified incubator at 37°C and 5% $CO_2$ under continuous gentle agitation to facilitate oxygen and nutrient exchange. Contractile force generation was monitored throughout culture, enabling real-time assessment of contractile activity and force development over time, analyzed with LabChart Reader 8.1.3. This biomimetic 3D culture platform served as the common experimental backbone for subsequent studies, including native myocardial slice culture, decellularized ECM-based constructs, and non-injury and injury model experiments. Experiment-specific interventions and readouts are described in the corresponding subsections.

**Method S15: Repopulation of decellularized myocardial ECM with DTPA-R HVPs**

Decellularized myocardial ECM scaffolds were generated from porcine left ventricular myocardium using an established detergent-based decellularization protocol optimized for cardiac tissue, as previously described (*7*). Briefly, myocardial tissue was incubated overnight at room temperature in lysis buffer containing 10 mM Tris and 0.1% EDTA (pH 7.4) under continuous agitation. The tissue was subsequently incubated in 0.5% SDS at room temperature for at least 6 h. After three washes with PBS, the resulting ECM scaffolds were incubated overnight at 4°C in 50% FBS prepared in PBS, followed by a final PBS wash. Prior to cell seeding, decellularized ECM scaffolds were equilibrated in culture medium. CTRL HVPs or DTPA-R HVPs were seeded onto ECM scaffolds at a density of approximately $2\times10^6$ cells per scaffold. Initial cell attachment was performed for 24 h in RPMI 1640 containing B-27 Supplement without insulin and 10 μM ROCK inhibitor Y-27632. Cultures were then maintained in RPMI 1640 containing B-27 Supplement without insulin until day 8, followed by transition to RPMI 1640 containing B-27 Supplement with insulin from day 8 onward. Cell-seeded ECM constructs were cultured for up to 14 days under biomimetic conditions using the chamber system as described. At experimental endpoints, constructs were fixed and processed for immunofluorescence staining of sarcomeric α-actinin and V5-tag, and nuclei were counterstained with Hoechst 33258.

**Method S16: Radiofrequency ablation injury**

RFA injury was generated using a THERMOCOOL SF unidirectional catheter with a 3.5-mm tip electrode (Biosense Webster) connected to a Stockert 70 radiofrequency generator. Radiofrequency energy was delivered at 20 W for 15 s, producing a localized, non-transmural lesion. Following focal cell seeding ($1\times10^6$) opposite the injury, slices were cultured under identical dynamic conditions. Two different cohorts were imaged separately at day 2 and day 5 post-injury after cell seeding.

**Method S17: [$^{18}$F]F-DTPA•Tb PET imaging myocardial slices**

Static PET acquisition was performed for 20 min using a small-animal PET/CT scanner (Inveon, Siemens Preclinical Solutions). Images were reconstructed using a three-dimensional ordered-subset expectation maximization (OSEM) algorithm and analyzed with Inveon Research Workplace (IRW, version 4.2). A predefined square region of interest encompassing the entire myocardial slice was applied, and decay-corrected activity concentration (Bq $mL^{-1}$) was quantified. Total reconstructed activity (Bq) was calculated by multiplying the measured activity concentration by the corresponding ROI volume.

**Method S18: Imaging with a clinical PET/MR scanner**

The 200 μL sample was sealed in a 200 μL pipette tip, and the 1,000 μL sample was sealed in a syringe barrel. Both samples were positioned side by side within a central region of the phantom to ensure identical attenuation, scatter, and imaging geometry. Static PET acquisitions (20 min) were acquired at 0, 2, 4, and 6 h after sample preparation. PET data were reconstructed using the manufacturer's standard workflow and analyzed with Inveon Research Workplace (Siemens Healthineers). Square regions of interest encompassing each hotspot were applied to quantify activity concentration ($Bq\cdot mL^{-1}$). To verify equal total activity, aliquots from each sample were independently measured in a gamma counter.

### Supplementary Tables

#### Table S1: List of antibodies and fluorescent probes

| Antibodies (IF) | Host / Clone | Source | Identifier | Dilution |
|---|---|---|---|---|
| Anti-V5-tag | Mouse monoclonal, SV5-Pk1 | Bio-Rad | MCA1360 | 1:250 (2D); 1:100 (3D) |
| Anti-V5-tag | Mouse monoclonal, SV5-Pk1 | Abcam | ab27671 | 1:500 (2D); 1:250 (3D) |
| Anti-NANOG | Rabbit polyclonal | Abcam | ab21624 | 1:500 |
| Anti-OCT4 | Rabbit polyclonal | Abcam | ab19857 | 1:200 |
| Anti-ISL1 | Mouse monoclonal, 39.4D5 | Hybridoma Bank | 39.4D5 | 1:100 |
| Anti-NKX2-5 | Rabbit polyclonal | Thermo Fisher | PA5-49431 | 1:200 |
| Anti-cardiac troponin T (cTNT) | Rabbit recombinant monoclonal, EPR3696 | Abcam | ab92546 | 1:500 (2D); 1:250 (3D) |
| Myosin Light Chain 2/MLC-2V | Rabbit polyclonal | Proteintech | 10906-1-AP | 1:100 |
| Sarcomeric anti-α-actinin | Mouse monoclonal | Sigma | A7811 | 1:300 |
| Alexa Fluor 488, goat anti-mouse | Polyclonal | Invitrogen Molecular Probes | A-11001 | 1:500 |
| Alexa Fluor 647, goat anti-mouse | Polyclonal | Invitrogen Molecular Probes | A-21235 | 1:500 |
| Alexa Fluor 488, goat anti-rabbit | Polyclonal | Invitrogen Molecular Probes | A-11008 | 1:500 |
| Alexa Fluor 647, goat anti-rabbit | Polyclonal | Invitrogen Molecular Probes | A-21244 | 1:500 |
| Hoechst 33258 | n.a. | Abcam | ab228550 | 5 µg $mL^{-1}$ |
| **Antibodies (flow cytometry)** | | | | |
| Anti-V5-tag, Alexa Fluor 488 | SV5-Pk1 | Bio-Rad | MCA1360A488 | 1:100 |
| Isotype Control Antibody, mouse IgG2a | S43.10 | Miltenyi Biotec | 130-113-271 | 1:100 |
| Anti-TRA-1-60, PE | REA157 | Miltenyi Biotec | 130-122-921 | 1:100 |
| REA Control Antibody (S), human IgG1, PE | REA293 | Miltenyi Biotec | 130-113-438 | 1:100 |
| Anti-cardiac troponin T (cTNT), APC | REA400 | Miltenyi Biotec | 130-120-403 | 1:100 |
| REA Control Antibody (I), human IgG1, APC | REA293 | Miltenyi Biotec | 130-120-709 | 1:100 |

### Supplementary Figures

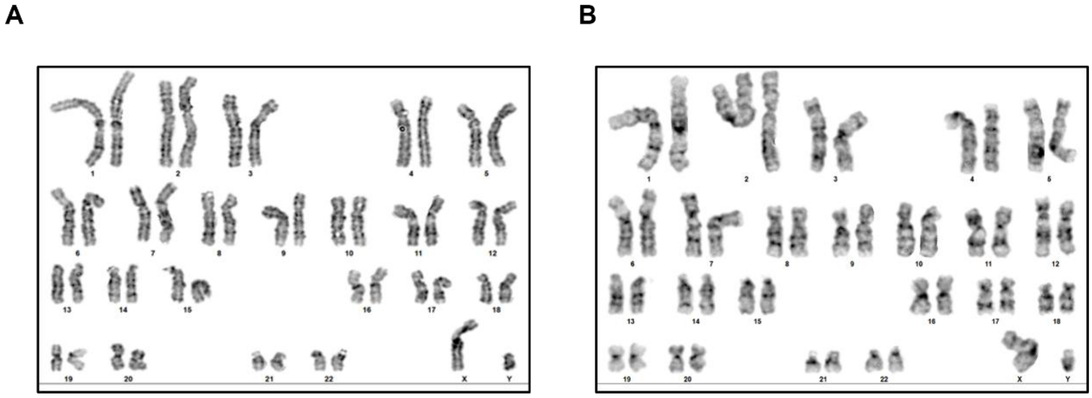

**Figure S1: G-banded karyotype analysis of CRISPR/Cas9-engineered DTPA-R hiPSC clones.** Conventional G-banding was performed to assess genomic integrity following targeted integration of the DTPA-R construct at the AAVS1 locus. **A,** Homozygous clone and **B,** heterozygous clone, both exhibiting a normal human karyotype without detectable structural or numerical chromosomal abnormalities.

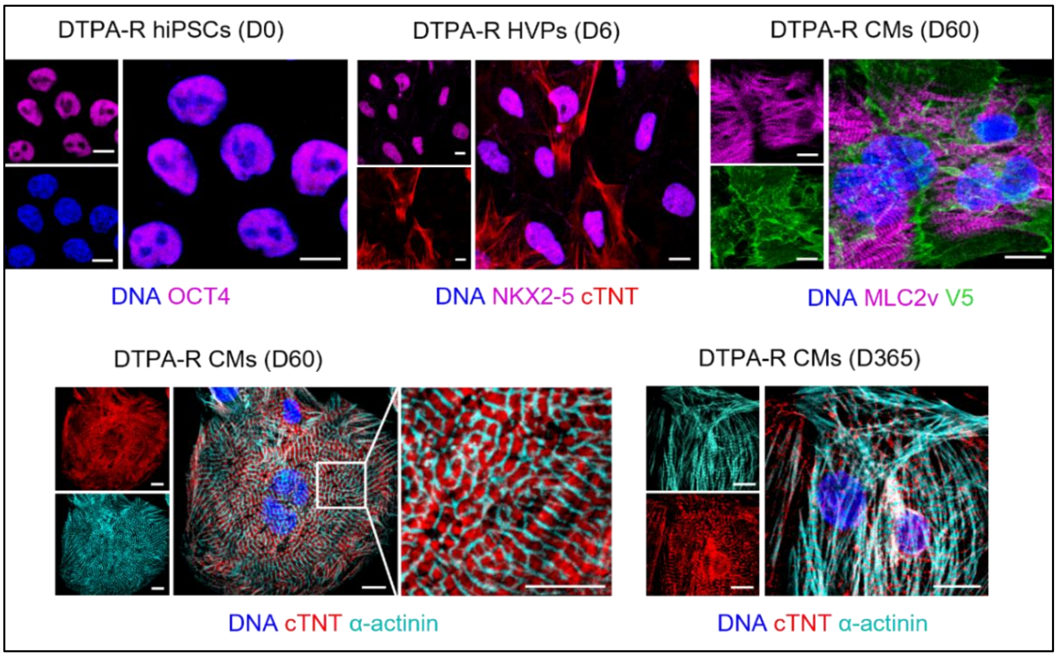

**Figure S2: Immunofluorescence analysis of DTPA-R hiPSCs and cardiac derivatives HVPs and CMs in 2D culture.** Representative immunofluorescence images of DTPA-R hiPSCs and derived cells across differentiation stages. hiPSCs (day 0) were stained for OCT4 (magenta), and HVPs (day 6) were stained for NKX2-5 (magenta) and cTNT (red). CMs (day 60) were stained for ventricular myosin light chain 2 (MLC2v, magenta), V5-tag (green), cTNT (red) and α-actinin (magenta), including higher-magnification visualization of sarcomeric organization. CMs (day 365) stained for cTNT (red) and α-actinin (magenta). Images are representative of three biological replicates (n=3).. Scale bars, 10 µm.

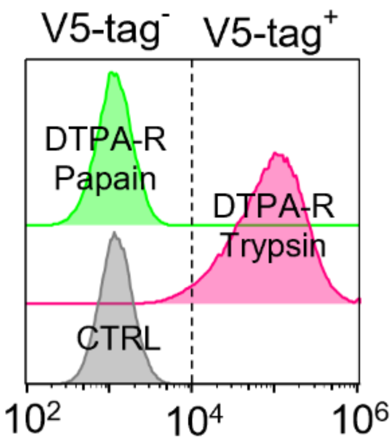

**Figure S3: Trypsin dissociation of HVPs preserves DTPA-R detection on cell surface.** Representative flow cytometry histograms acquired immediately after enzymatic dissociation of DTPA-R HVPs. Trypsin-treated cells (magenta) retain a clearly detectable V5-tag positive population compared to control cells (gray), whereas papain-treated cells (green) overlap with the V5-tag negative control distribution. The dashed vertical line indicates the gating threshold separating V5-tag–negative (left) and V5-tag–positive (right) populations. These data demonstrate that trypsin preserves a detectable surface V5-tag signal, while papain reduces it to background levels.

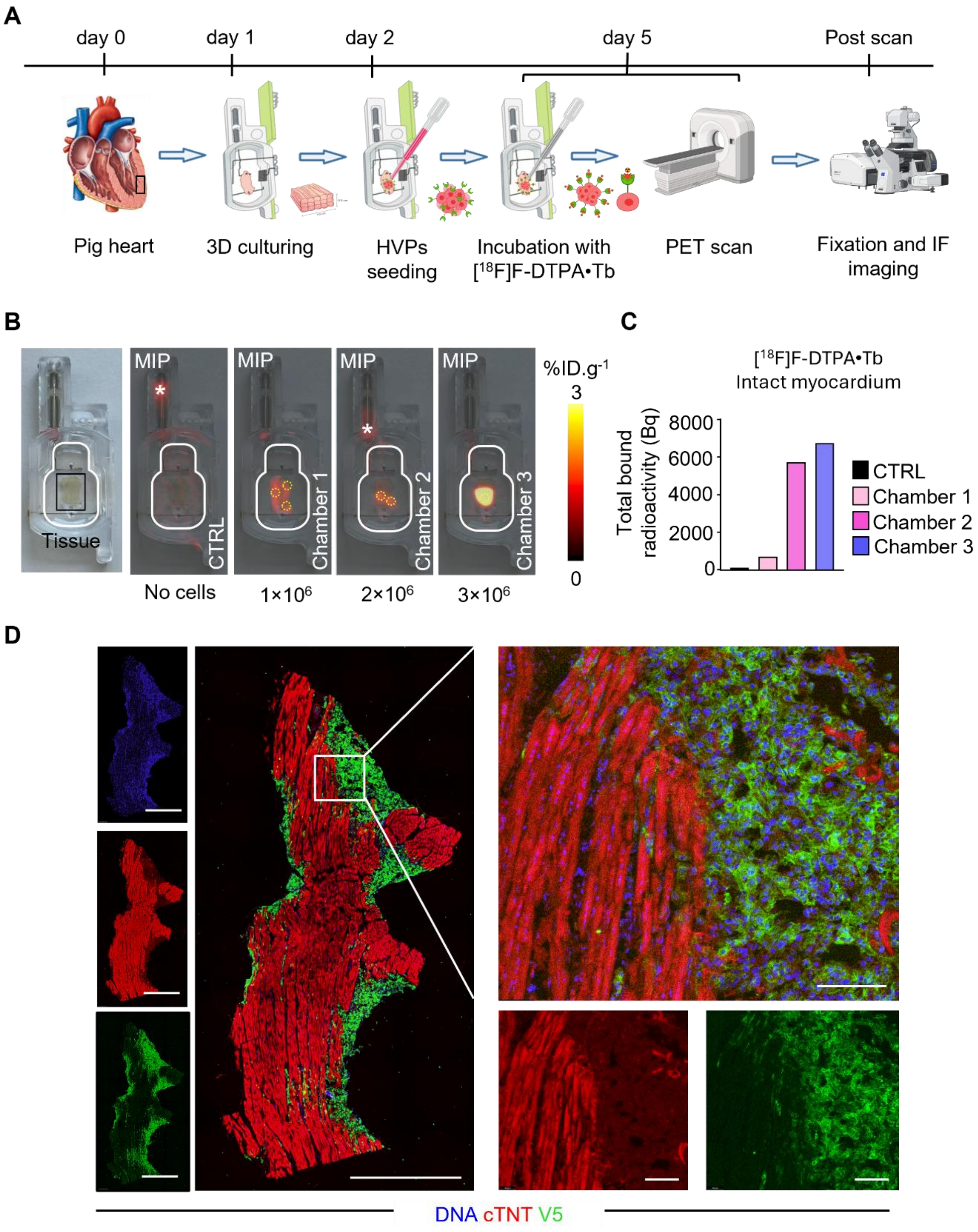

**Figure S4: *Ex vivo* PET detection of DTPA-R HVPs in native pig heart slices. A,** Schematic of the *ex vivo* heart slice imaging workflow. Native porcine ventricular slices were cultured in a biomimetic chamber, locally seeded with different numbers of DTPA-R HVPs, radiolabeled with [18F]F-DTPA•Tb 3 days after seeding, imaged by PET, and subsequently processed for immunofluorescence. **B,** Maximum intensity projection (MIP) PET images of native pig heart slices cultured without cells (CTRL) or seeded with DTPA-R HVPs under non-injury conditions. Seeding configurations included $1\times10^6$ cells distributed across three sites (chamber 1), $2\times10^6$ cells across two sites (chamber 2), or $3\times10^6$ cells at a single site (chamber 3). PET signal intensity is displayed as %ID·g$^{-1}$. Asterisks indicate nonspecific radiotracer accumulation outside of the chambers. Dashed white circles outline the cell seeding sites. **C,** Quantification of total PET-detected radioactivity within tissue volumes of interest under the indicated seeding conditions, expressed as Bq. **D,** Representative immunofluorescence tile-scan images of heart slices following PET imaging and fixation, stained for cTNT (red), V5-tag (green) and Hoechst 33258 (blue). Left panels show individual channels and merged tile-scan overviews; right panels show higher-magnification views of boxed regions. Scale bars, 1 mm (overviews) and 100 µm (magnified views).

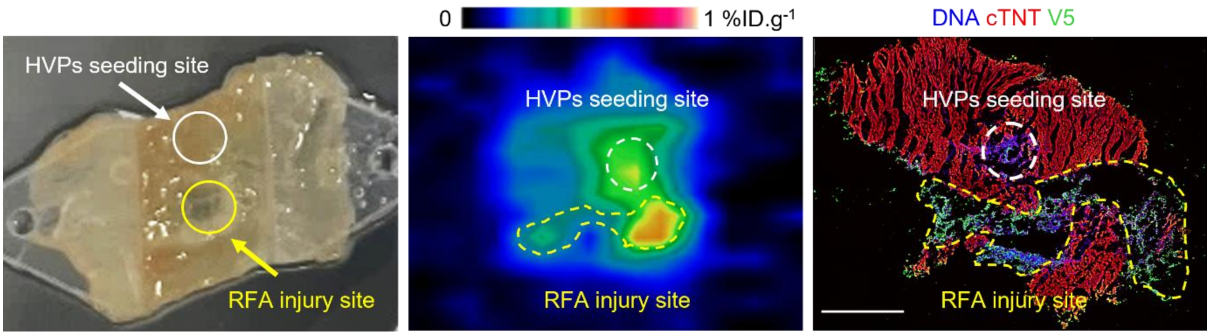

**Figure S5: PET signal detected in the RFA injury site on day 5 post-injury**. Representative images of an *ex vivo* porcine myocardial slice at day 5 following HVP seeding and RFA injury. Left, bright-field image of the myocardial slice indicating the HVP seeding site (white circle) and the RFA injury site (yellow circle). Middle, corresponding PET image demonstrating detectable signal at both sites, with higher signal intensity in the RFA injury region. Right, corresponding merged immunofluorescence image stained for cTNT (red), V5-tag (green) and Hoechst 33258 (blue). Dashed circles indicate the HVP seeding site (white) and the RFA injury site (yellow) at the selected plane. Scale bar (immunofluorescence), 1 mm.

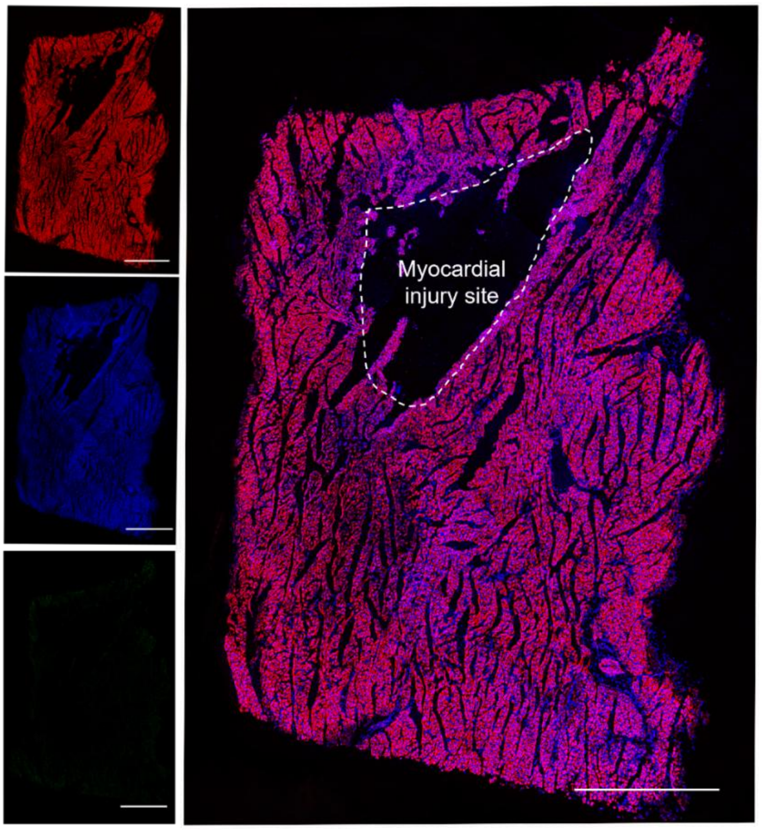

**Figure S6: Immunofluorescence analysis of *ex vivo* control RFA-injured myocardial slice**. Representative immunofluorescence images of *ex vivo* porcine myocardial slices at day 2 following RFA injury without application of therapeutic cells. Representative images of cTNT (red), V5-tag (green), and Hoechst 33258 (blue). Left, individual channels; right, merged tile scan. The scale bar represents 1 mm. The RFA-induced lesion area, outlined by a dashed line, exhibits reduced and disrupted cTNT and Hoechst signal compared to the surrounding non-injured myocardium.
